# Spec2Vec: Improved mass spectral similarity scoring through learning of structural relationships

**DOI:** 10.1101/2020.08.11.245928

**Authors:** Florian Huber, Lars Ridder, Stefan Verhoeven, Jurriaan H. Spaaks, Faruk Diblen, Simon Rogers, Justin J.J. van der Hooft

**Affiliations:** Netherlands eScience Center, Amsterdam, the Netherlands; School of Computing Science, University of Glasgow, Glasgow, United Kingdom; Bioinformatics Group, Plant Sciences Group, University of Wageningen, Wageningen, the Netherlands

## Abstract

Spectral similarity is used as a proxy for structural similarity in many tandem mass spectrometry (MS/MS) based metabolomics analyses such as library matching and molecular networking. Although weaknesses in the relationship between spectral similarity scores and the true structural similarities have been described, little development of alternative scores has been undertaken. Here, we introduce Spec2Vec, a novel spectral similarity score inspired by a natural language processing algorithm -- Word2Vec. Spec2Vec learns fragmental relationships within a large set of spectral data to derive abstract spectral embeddings that can be used to assess spectral similarities. Using data derived from GNPS MS/MS libraries including spectra for nearly 13,000 unique molecules, we show how Spec2Vec scores correlate better with structural similarity than cosine-based scores. We demonstrate the advantages of Spec2Vec in library matching and molecular networking. Spec2Vec is computationally more scalable allowing structural analogue searches in large databases within seconds.

## Introduction

In metabolomics the high throughput characterisation of metabolites present in a biological sample, is increasingly important across the biomedical and life sciences ^1,2^. This is largely due to the manner in which the metabolome complements the genome, transcriptome and proteome as the data type most closely representing phenotype^3^, and to the increased sensitivity and coverage of modern measurement equipment.

Of the available measurement platforms, liquid chromatography coupled to mass spectrometry (LC/MS) is the most widely used. Modern untargeted LC/MS experiments produce large datasets that are challenging to fully analyse and exploit. A main bottleneck is the structural annotation and identification of the chemical ions detected by the mass spectrometer, primarily because the mass-to-charge ratio (m/z) of an observed ion species is very often insufficient to unambiguously assign it to one chemical formula, and certainly insufficient to assign it to a specific chemical structure. Building upon the assumption that molecules fragment in a manner that is dependent on their structure, fragment data (also known as MS2 or MS/MS) is often used to overcome this bottleneck. Fragmentation spectra can act as an aid to annotation via either comparison with databases ^4^ such as MassBank^5^, Metlin^6^, GNPS^7^, etc, or as input to in-silico identification algorithms such as SIRIUS/CSI:Finger ID ^8^ or MS2LDA ^9^.

At the heart of much analysis of mass spectrometry fragmentation data is the computation of similarity between pairs of MS2 spectra (Fig.1 A,B). For example, when searching an unknown spectrum against a database, a similarity score is computed between the query spectrum and database spectra. Similarly, when creating molecular networks ^10^, edges are drawn between spectra if their similarity exceeds a user-defined threshold. In all such analyses, spectral similarity is being used as a proxy for structural similarity, the real quantity of interest ^11^. Cosine-based scores are the most widely used measures of spectral similarity. Studies investigating whether molecules with high structural similarity result in spectra with a high spectral cosine similarity score only partly support the assumed relationship between spectral and structural similarity ^12,13^. As a result, various modifications of the cosine similarity score have been proposed, including raising the m/z and intensity components to different powers, and shifting fragment peaks by the difference in precursor m/z (‘modified cosine score’ ^10^). Cosine-based methods are very good at revealing nearly equal spectra, but by design they are not well-suited to handle molecules with multiple local chemical modifications. In addition, most cosine scores are computed by aligning the fragment peaks from the two spectra, a computationally intensive procedure that makes extensive library searching slow. Despite these limitations, thus far no fundamentally different spectral similarity scores have been proposed.

**Figure 1.**
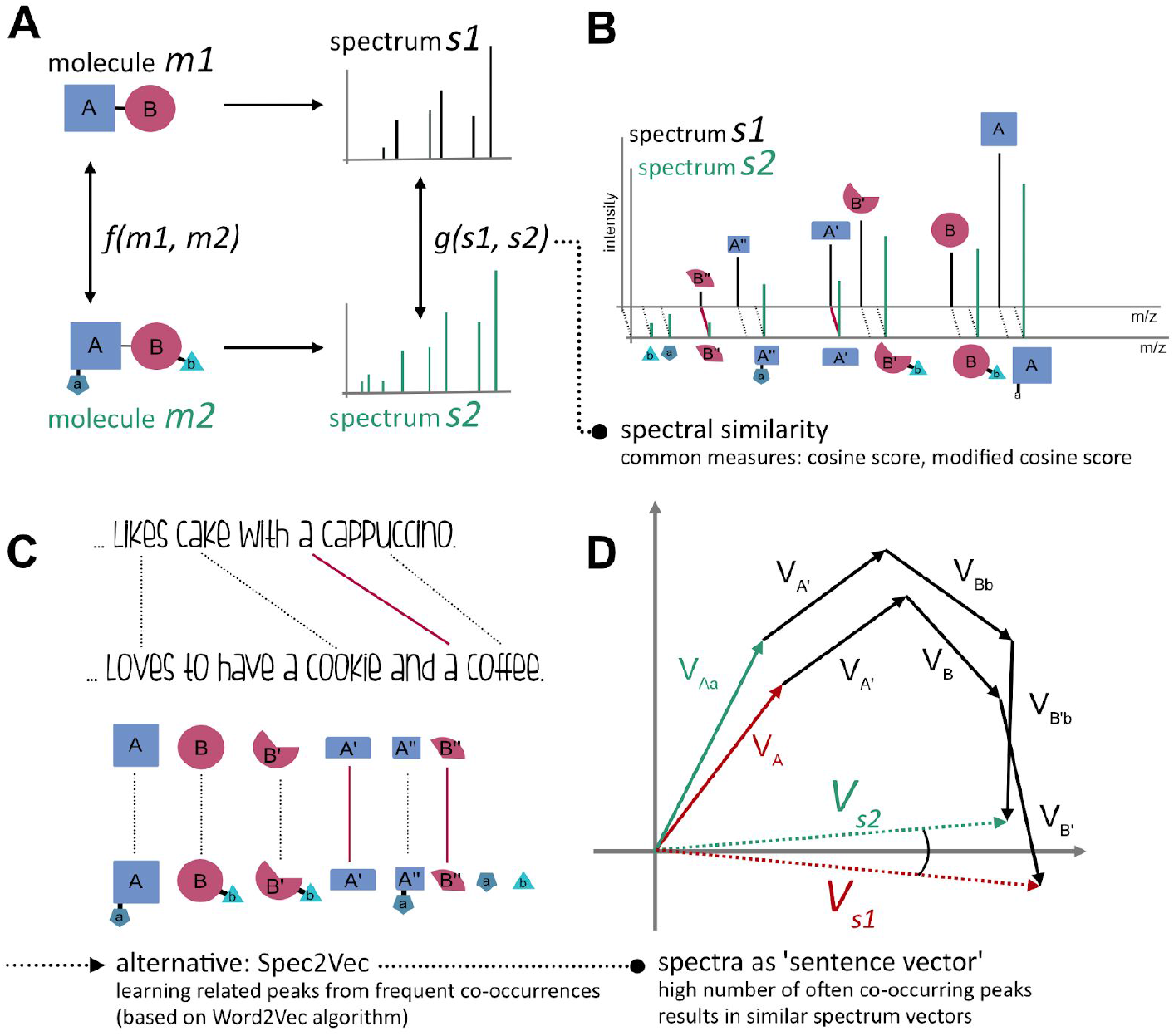
(A) MS-MS spectra can be considered as signatures of molecules: spectra are known to contain structural information of the original molecule, but without a straightforward way to translate mass spectral features into structural ones describing the fragmented molecule. (B) Spectra are commonly compared by similarity measures such as cosine or modified cosine scores. While those measures are very good at revealing (nearly-) equal spectra, they often underperform when it comes to spectra of complex molecules with high structural similarity, but which differ in multiple locations (C) Spec2Vec is based on algorithms from natural language processing and learns relationships between peaks based on how frequently they co-occur. (D) Two spectra from different yet similar molecules will hence be represented by similar spectral vectors even if many of their peak positions will differ.

Within the context of metabolite identification from MS/MS data, much effort has gone into the prediction of structural information from spectra. This is because it allows spectra to be queried against structural databases that are typically orders of magnitude larger than spectral ones. Despite the fact that database searching is still the gold standard for metabolite annotation^14^, methods such as MAGMa^15^, SIRIUS^16^, CSI:FingerID^8^, IOKR^17^, DeepMASS^18^, and MetDNA^19^ have been effective at widening the search space and clearly demonstrate that useful structural information can be learnt from MS2 spectra.

Building on that insight, we present a novel spectral similarity score based upon learnt embeddings of spectra. Inspired by the success of algorithms from the field of natural language processing in accounting for element relatedness for overall similarity assessment of objects, we aimed at adapting such tools to mass spectra data. A language-based analogy would be words like ‘cookie’ and ‘cake’ which often occur in similar contexts (e.g. together with words like ‘dough’, ‘sweet’, ‘eating’) and are hence assumed by the model to represent related entities (Fig.1 C).

Adapting Word2Vec, a well-established machine learning technique in natural language processing^20^, Spec2Vec learns from co-occurrences across large datasets to represent highly related fragments or neutral losses by vectors pointing in similar directions within a continuous abstract space. A spectrum can then be represented by a low-dimensional vector calculated as the weighted sum of all its fragment (and loss) vectors (Fig.1 D). Instead of relying on only a binary assessment of each fragment (match/no match), Spec2Vec hence takes the relation between fragments into account. This can be illustrated by comparing two spectra of molecules with multiple (small) chemical modifications that Spec2Vec could relate to each other despite the low amount of direct peak matches (Fig. 2).

**Figure 2.**
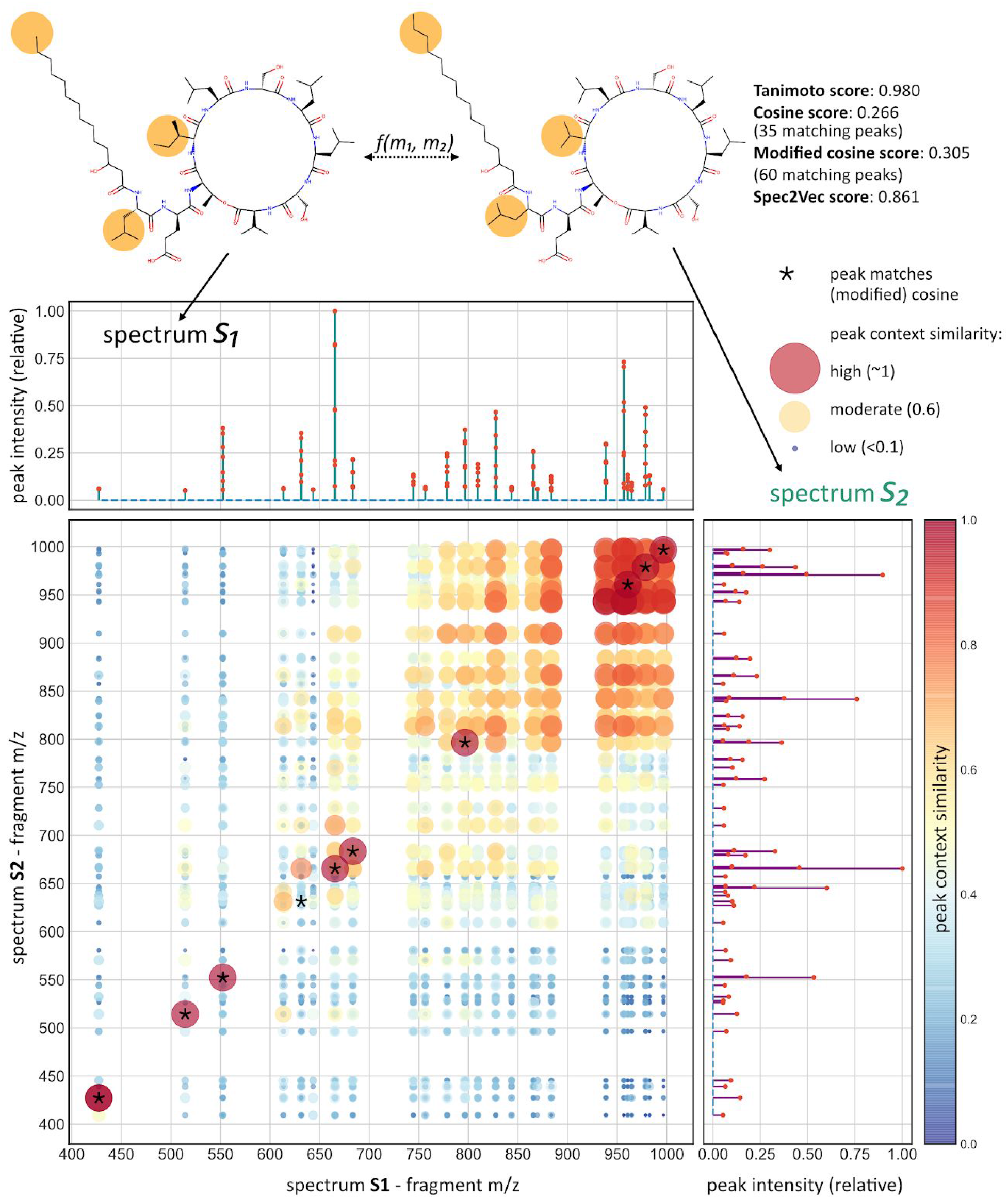
In-depth comparison example of two spectra. Since the two molecules differ slightly in three locations, both cosine and modified cosine scores fail to recognize the overall structural similarity and return low spectral similarity scores. Spec2vec for many peaks acknowledges that they often co-occur across the training data, hence showing a high peak context similarity which overall leads to a high Spec2Vec similarity score. For illustrative purposes, this figure only displays peaks between 400 and 1000 Da.

In contrast to the database searching methods mentioned above, Spec2Vec is unsupervised and can be trained on any collection of spectra. We demonstrate that Spec2Vec similarity scores are well suited to identify structural similarities on the basis of given MS/MS spectra. Spec2Vec similarity scores are very fast to compute, which -taken together-creates promising use cases of Spec2Vec in library matching as well as in molecular networking.

## Results

### Spectral similarity vs structural similarity

In order to quantify how well different spectral similarity scores correlate with structural similarity, we calculated multiple spectral similarity scores for all possible pairs across a mass spectra dataset of representative character, but computationally manageable size. To create this dataset, we started with a large collection of mass spectra provided by GNPS (see Methods). Library spectra were used to compare spectral similarity scores with scores based on (the known) molecular structures. It is important to note, however, that Spec2Vec is an unsupervised machine learning technique that can be trained on any collection of spectra, independent of whether the chemical structures are known. After filtering, processing, and removing all spectra with fewer than 10 fragment peaks (see Methods), the remaining **AllPositive** dataset comprised 95,320 positive ionization mode mass spectra, 77,092 of which had InChIKey annotations. Because many similarity scores are computationally expensive, the quantitative similarity score assessment was done on a subset of this data, **UniqueInchikey,** consisting of 12,797 spectra with unique InChIKeys (first 14 characters, also termed planar InChIKeys, see Methods).

For the **UniqueInchikey** data it was possible to compare the different spectra similarity scores to the structural similarity, represented by Tanimoto scores. One of our core interests was to evaluate to what extent a high spectral similarity score, *g*(*s*_1_,*s*_2_) between a pair of spectra reflects a high structural similarity score *f*(*m*_1_, *m*_2_) between the respective molecules. For this, we computed different similarity scores for all possible spectra pairs, hence between all 12,797 spectra in **UniqueInchikey**. The vast majority of those spectra pairs correspond to entirely unrelated molecules, resulting in a distribution of fingerprint-based structural similarities as shown in Fig. 3A. We then selected only the structural similarities that correspond to the highest scoring pairs according to each similarity measure (Spec2Vec, Cosine, modified Cosine). Figure 3B displays the average structural similarity over the highest 0.1% of each respective spectra similarity score, with 0.1% corresponding to about 80,000 spectra pairs. This reveals that a high Spec2Vec spectrum similarity score correlates stronger with structural similarity than the cosine or modified cosine scores (Fig.3). As a consequence, Spec2Vec similarities allow retrieving notable larger fractions of spectra pairs above a desired mean structural similarity score (see example in Fig.3 B). Cosine scores exist in numerous flavors (e.g. using different peak weighting) and can vary largely depending on their key parameters (tolerance and min_match, the minimum number of matching peaks). Several different cosine score flavors and parameter ranges were tested, without resulting in major improvements regarding figure 3, see also supplemental material and figures S2 to S6.

**Figure 3.**
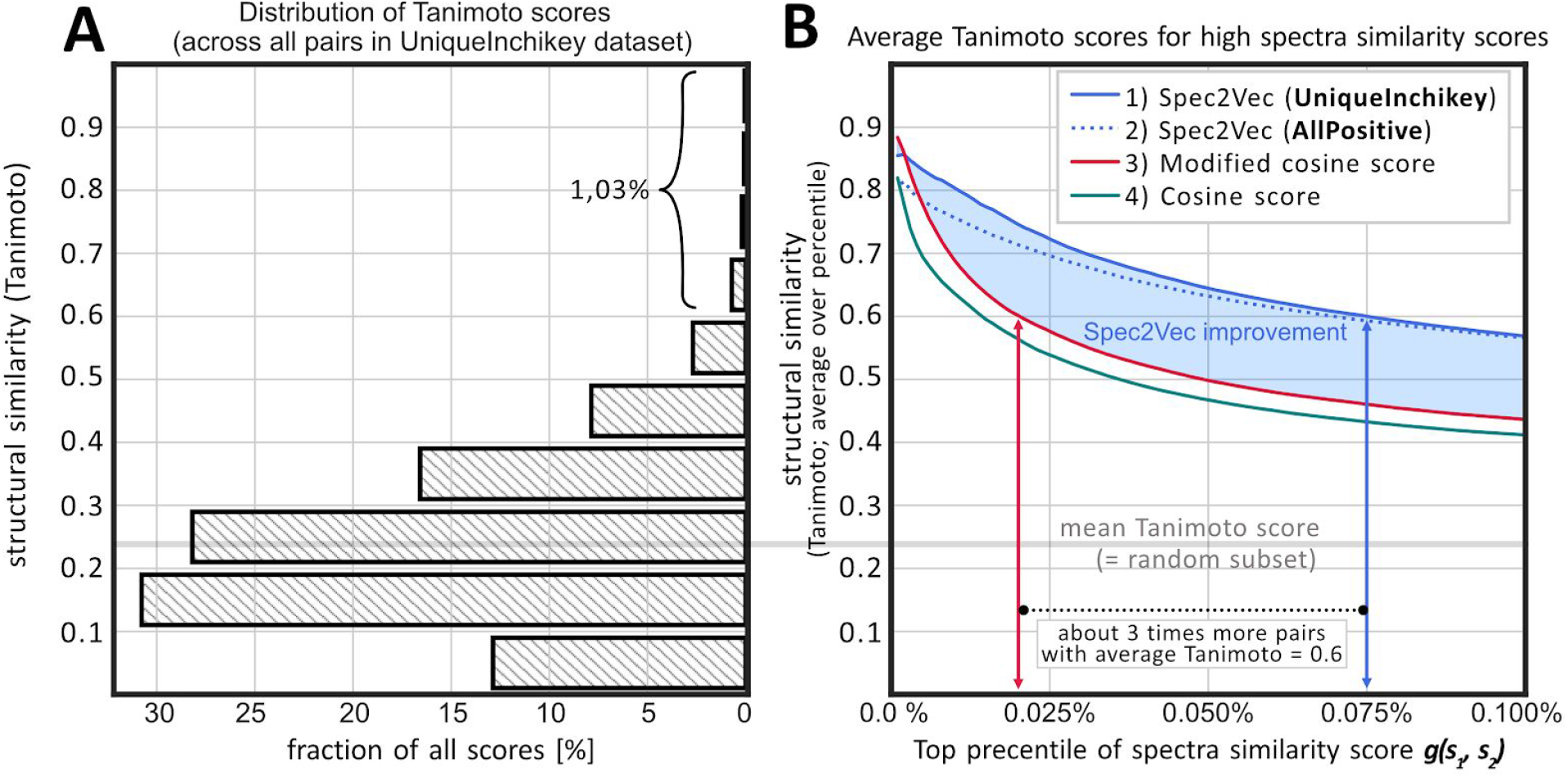
(A) histogram of the structural similarity scores across all possible spectra pairs between the 12,797 spectra in the **UniqueInchikey** dataset (81,875,206 unique pairs, not including pairs of spectra with themselves). The histogram indicates that randomly chosen pairs will most likely show scores between 0 and 0.5. Similarity scores > 0.6 are rare and hence unlikely to achieve by randomly choosing pairs (p=0.0103 of finding a score > 0.6, p=0.0034 for a score > 0.7). (B) Different similarity scores were calculated for the same 81,875,206 spectral pairs. Comparing the highest 0.1% the resulting scores to the structural similarities reveals that Spec2Vec similarities show a notably higher correlation with actual structural similarities. Used parameters were 1) Spec2Vec, trained on UniqueInchikey for 50 iterations, 2) Spec2Vec, trained on AllPositive for 15 iterations, 3) Modified cosine score with tolerance=0.005 and min_match=10, and 4) Cosine score with tolerance=0.005 and min_match=6.

We observed that the poorer correlation between cosine and modified cosine similarity scores and structural similarity can largely be explained by high false positive rates (Fig. S1). This shortcoming can to some extent -- though never fully -- be reduced by using lower peak match tolerances (here: tolerance = 0.005 Da) and ignoring scores based on fewer than *min_match* matching peaks (here: *min_match*=10, see Fig. S2, S3 for parameter search).

### Library matching

Next, we evaluated the potential of Spec2Vec to aid in matching unknown spectra to library spectra run on various instruments under different conditions. We worked with the **AllPositive** dataset (95,320 spectra), which we split into a library set (94,320 spectra) and a query set (1000 spectra). The query spectra were randomly selected such that they would all have a different planar InChIKey and that we had at least 4 spectra with identical InChIKey remaining in the library set. Therefore, for each of the 1000 query spectra, multiple positive hits existed in the library set. A Spec2Vec model was trained on the library set and Spec2Vec scores were compared with cosine similarity scores for library matching (Fig.4). Both Spec2Vec and cosine similarity scores were used in the same way: potentially matching spectra were pre-selected based on precursor mass matches (tolerance = 1Da) before the highest scoring candidate above a similarity threshold was chosen. Gradually lowering this threshold from 0.95 to 0, increases both the number of true and false positives per query. While the general trend for both scores is similar, Spec2Vec resulted in a notably better true/false positive ratio at all thresholds. Spec2Vec also allowed to correctly match the query spectra with up to 87% accuracy and showed both higher accuracy and retrieval rates when compared to the cosine score based library matching (Fig. 4).

**Figure 4.**
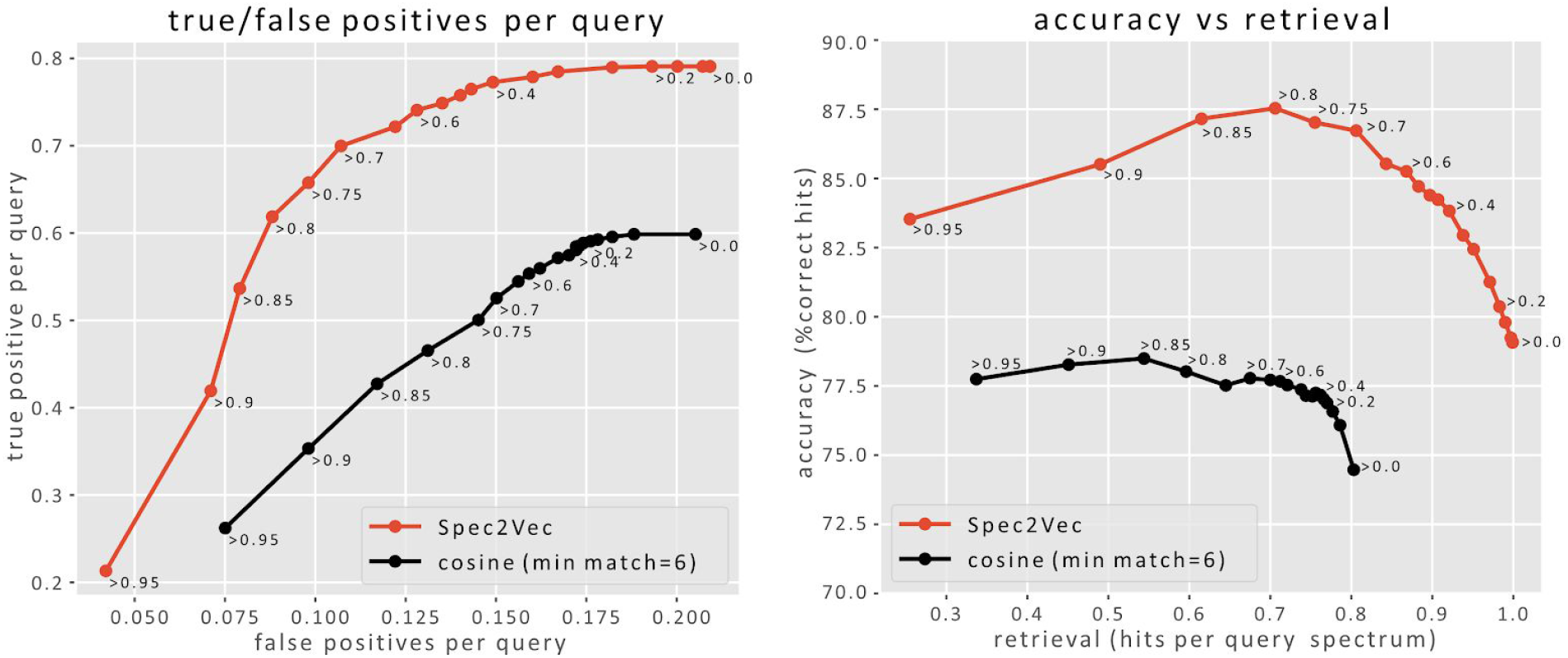
Spec2Vec similarity scores deliver improved true-to-false-positive ratios during library matching. 1000 randomly selected spectra, all with at least 5 identical InChIKeys in the entire dataset, were removed from a **AllPositive** and then matched to the remaining spectra. Matching was done by pre-selecting spectra with the same parent mass (tolerance = 1Da) and then choosing the candidate with the highest spectral similarity score if this score was larger than a set threshold. The left plot shows the true-vs-false ratio per query when using Spec2Vec (red) or Cosine scores (black). Labels near the dots report the used similarity score thresholds. The plot on the right displays the resulting accuracy and retrieval rates for the same parameters. Using Spec2Vec, library matching could be done with notably higher accuracy across all tested retrieval rates.

In practice, we expect that actual library matching can be improved further via stricter parent mass matching (lower tolerance) or by consulting both Spec2Vec and Cosine similarity scores, which can potentially make use of the apparent complementarity of the methods which is described in the next section.

### Unknown compound matching

Once the embedding has been trained, Spec2Vec similarity scores are computationally more efficient to calculate than cosine-based scores, allowing brute-force all-vs-all comparisons of query spectra against large datasets. Although computation times for the different similarity scores will depend upon multiple factors such as the chosen spectra preprocessing steps (key variable: resulting number of peaks), and implementation or hardware details, we generally found Spec2Vec similarities to be about 2 orders of magnitude faster to calculate when using a pre-trained model, and about 1 order of magnitude faster when training a new model from scratch. Training a model on the **UniqueInchikey** dataset takes about 30 minutes on a Intel i7-8550U CPU. The main performance gain results from the fact that Spec2Vec embedding vectors are fixed length vectors, comparison of which is straightforward. Conversely, cosine-based methods, despite their name, do not tend to operate on fixed lengths vectors as this would require a binning step. Instead, they rely on a costly alignment step in which all pairs of fragment ions with matching m/z (within some tolerance) are extracted. In addition, Spec2Vec similarity correlates more closely with structural similarity (Fig.3) which makes it better suited for identifying relationships between spectra of different yet chemically related molecules. Taken together, this makes it possible to use Spec2Vec similarity for rapidly querying spectra of unknown molecules against all spectra in a large database (Fig.5).

**Figure 5.**
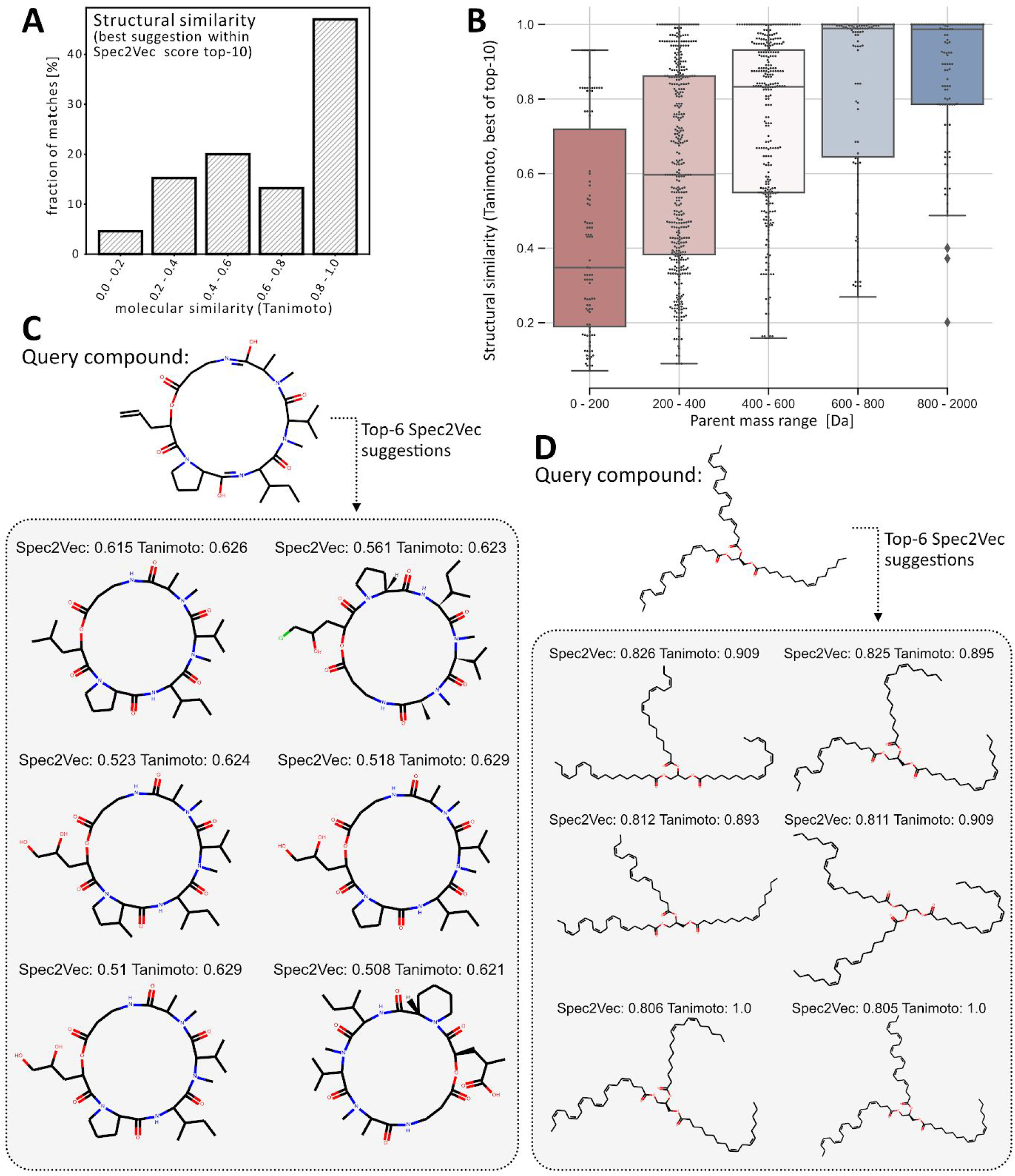
Matching of unknown molecules (not part of library) using Spec2vecs similarities. All spectra of 200 randomly selected InChIkeys (1030 spectra) were removed from the **AllPositive** dataset. Using a word2vec model that was trained on the remaining dataset, also excluding non-annotated spectra (76,062 spectra), each removed “query” spectrum was compared to the dataset by only using the Spec2Vec similarity score. (A) shows a histogram of the best structural similarity score out of the found top-10 Spec2Vec similarities for each query. For nearly 6 out of 10 queries, Spec2Vec finds a match with a structural similarity score > 0.6 reflecting high molecular similarity. (B) The quality of the suggested matches is highly dependent on the mass of the query compound. In particular for larger molecules (> 400 Da), Spec2Vec similarities allow finding highly similar molecules. (C+D) Examples of unknown molecules (not part of library) that are compared to all library spectra to find most similar matches using Spec2Vec. In both cases the algorithm is able to return highly related molecules to the query molecules that could be used to help with annotating the query spectra or to infer its chemical class.

To test whether Spec2Vec similarity scores can be used to help detect highly related molecules, we moved all spectra belonging to 200 randomly chosen InChIKey (1030 spectra in total) from the **AllPositive** dataset to a query dataset. To make sure that there is no overlap regarding molecules between training and query data, we also removed all spectra without InChIKey annotation. A separate Word2Vec model was trained on the remaining data of 76,062 spectra, and therefore the model did not see any spectra of the selected 200 molecules. Finally, each of the 1030 query spectra was compared to all remaining spectra. The ten highest scoring matches for each query were selected and the quality of this selection was assessed by computing the structural similarities between all found matches and the query molecules (Fig.5). For 3 out of 5 queries, the resulting top-10 list would contain suggested molecules with a structural similarity score of > 0.6 (p=0.0103, see Fig. 3A for the histogram of scores and calculation of the percentage). In particular for molecules with parent masses larger than 200 Da, Spec2Vec was often able to detect highly related molecules for unknown queries. For query spectra of larger molecules (>400 Da), Spec2Vec similarities were very likely to point to chemically related library molecules with an average Tanimoto similarity for the best suggestions above 0.8 (Fig.5 B).

This experiment purposely considered Spec2Vec as the only means to find related matches to best illustrate its ability to quickly find related molecules in a large library dataset. Querying 1030 spectra against 76,062 spectra took 110s on a Intel i7-8550U CPU, or 0.1s per query spectrum. This makes Spec2Vec similarity scores a powerful selection tool. With a total of 1, 030 · 76, 062 ≈ 78 · 10^6^ similarity scores to calculate, we estimate the same operation using cosine or modified cosine would take about 4-8 hours, which is the time it took to compute the 82 · 10^6^ similarity scores for the benchmarking comparison (Fig. 3). The high computational cost of the cosine and modified cosine scores is also the reason why potential matches are often pre-selected, for instance by comparing the parent masses. Spec2Vec in contrast, easily allows to compare large numbers of spectra purely based on their spectral similarity. In addition to the 2 orders of magnitude gain in computation time, our results also suggest that Spec2Vec is a better suited similarity score for detecting related yet different molecules. This can be seen in the considerably lower correlation between high spectral similarity and molecular similarity for the cosine and modified cosine score (Fig.3), as well as the observed high fraction of false positives (Figure S1).

Unlike in our controlled experiments, we will in practice generally not know if our library contains exact compound matches for our query spectra. We could expect the most reliable library search results when combining different measures. Thus, in the future, we could imagine that Spec2Vec similarity scores together with parent mass matching are very suitable methods for pre-selecting promising candidates for library matching. In a second step, computationally more expensive similarity measures, including cosine and modified cosine scores, but also scores such as those derived from SIRIUS/CSI:Finger ID^8^ could then be added to the Spec2Vec similarity measure to facilitate a well-informed decision. For instance, high Spec2Vec similarity combined with low cosine score could be used as a signature of a chemically related yet distinct compound. Having both a high Spec2vec and cosine score together with a parent mass match would then suggest an exact match.

### Network analysis

Based on the finding that Spec2Vec similarity correlates well with structural similarity (Fig.3), we investigated how Spec2Vec could be applied to molecular networking which is becoming an increasingly popular tool for exploring metabolomic datasets ^7^. Molecular networking refers to representing spectra as a network in which spectra are nodes connected by edges based upon a user defined cutoff for the similarity score (along with some heuristic pruning). Current molecular networking relies on the modified cosine score ^7^’^10^.

Such networks are frequently used to define clusters (or communities) which are termed molecular families. Detecting clusters in complex networks without ground truth is generally regarded as an ill-defined problem ^21^, which makes absolute quantitative comparisons difficult. In the present case, clusters depend strongly on the chosen parameters (e.g. similarity cutoff) and algorithm (processing, cleaning and trimming of the network). To better isolate the effect of different similarity scores we have chosen a simple workflow. For each spectrum (=node) the up to 10 highest-scoring links (=edges) to other spectra are added if those links have similarity scores above a given threshold. To improve the overall quality of the clustering, but also to make the results more robust we further apply the Louvain algorithm to split up large, poorly connected clusters ^21,22^. Graphs in figure 6 were generated using different similarity thresholds and the resulting clusters were quantified by counting them as well-clustered if the mean structural similarity within all cluster edges was ≥ 0.5, and poorly-clustered if it was < 0.5. The rationale behind setting the cutoff to 0.5 was the low probability of finding scores above 0.5 by chance as observed in the histogram of all possible structural similarity scores within the **UniqueInchikey** dataset (Fig. 3A). The results in fig. 6 show that Spec2Vec is able to cluster higher fractions of spectra into high structural similarity clusters when compared to the modified cosine. In terms of computation time, the presented molecular networking procedure requires to calculate the spectral similarities between all possible spectra pairs (≈82 · 10^6^ unique pairs), which is the same as the before described benchmarking. Spec2Vec similarity scores here reduced the computation time by about 2 orders of magnitude. Finally, we expect that Spec2Vec similarity scores bring new options for further improving molecular networking. A simple first proof-of-principle test on combining Spec2Vec similarity and modified cosines scores reveals that combined scores will likely be able to further increase clustering accuracy (see fig. S9). More extensive future work will be necessary to systematically explore the full potential of such score combinations.

**Figure 6.**
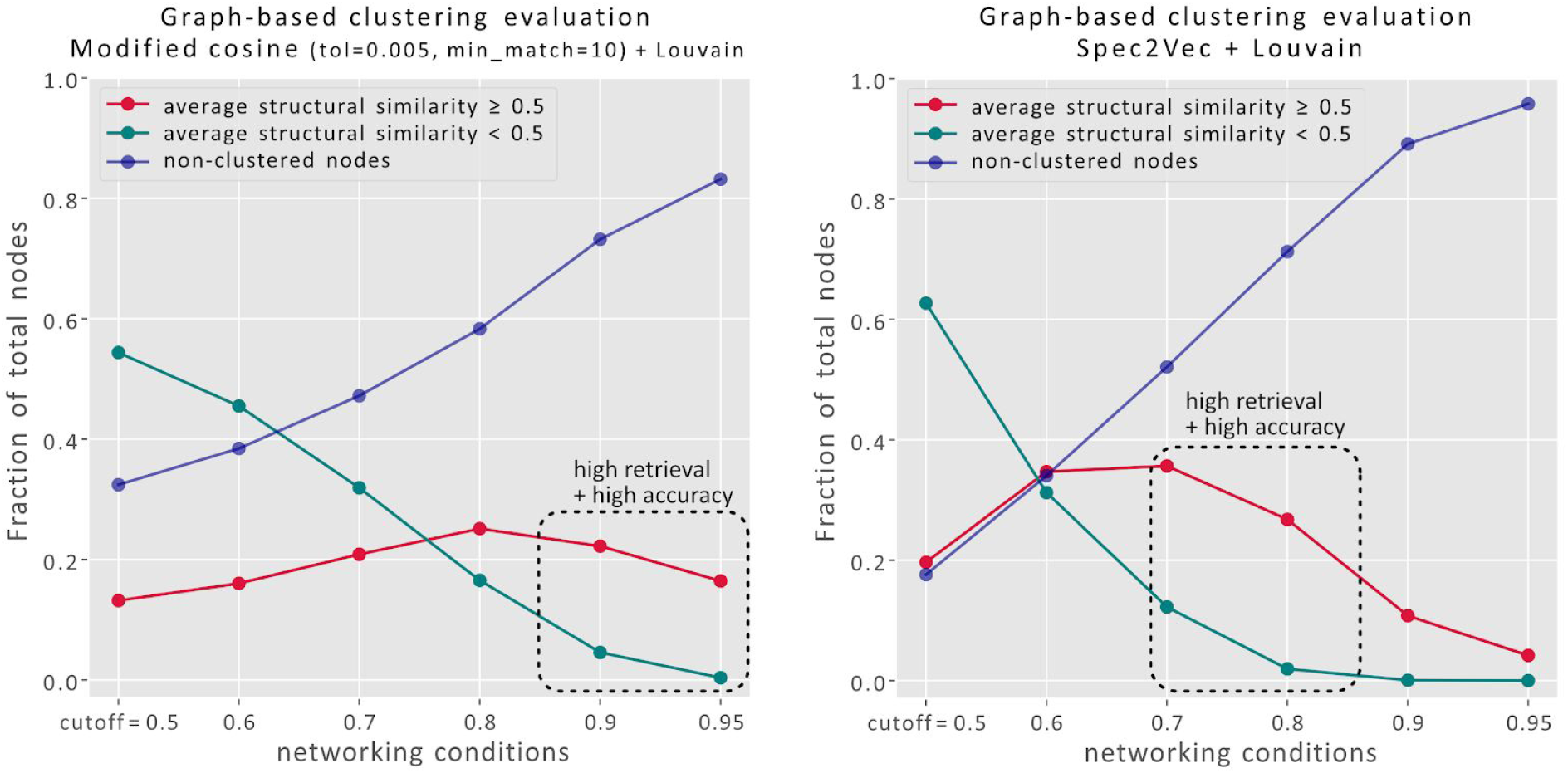
The better correlation between Spec2Vec spectral similarity with the actual structural similarity also translates into a more coherent clustering ability. Clustering is done here by creating edges between spectra (=nodes) for similarities above a certain cutoff (max. 10 links per node). To make the resulting clustering more robust and better comparable across different scores, we used the Louvain algorithm to break up the large clusters. Cluster quality here is assessed by measuring the average structural similarity across all linked pairs within each cluster. Setting a structural similarity threshold of 0.5 (see fig. 3A) allows to compare the fraction of spectra that ends up in chemically homogenous clusters (red) with those in more heterogeneous clusters (green) and the fraction of spectra that is not clustered at all (those with no links above set threshold). Dashed squares mark regions of relatively high retrieval (high fraction of clusters with high structural similarity) and high accuracy (large discrepancy between fraction of high structural similarity and low structural similarity clusters). Overall, Spec2Vec allows to cluster higher fractions of spectra into high structural similarity clusters (> 35% of all spectra are in high similarity clusters for a Spec2Vec similarity threshold of 0.7).

## Discussion

In conclusion, here we introduce a spectral similarity score with advantageous properties over the currently widely used cosine similarity score (and its popular variant, modified cosine). Inspired by natural language processing, much like topic modelling enabled substructure finding in metabolomics data^23^, we here show how a popular text mining algorithm (Word2Vec^20^) can be adapted to learn meaningful relations between mass fragments and neutral losses in mass fragmentation spectra. We demonstrate how the Spec2Vec score better resembles the structural similarity of fragmented molecules and outperforms the cosine score in key tasks underpinning metabolomics analyses including library matching. Being a machine learning algorithm, one limitation of Spec2Vec as compared to cosine scores is that it needs training data to learn the fragment peak relationships; however, since this not necessarily needs to be library spectra and in light of the enormous increase in publicly available metabolomics data, we do not see this as a major bottleneck. Our use of library data in this study was purely so that spectral similarities could be compared with the similarities of the known structures. Training time is not a limitation: training the embedding on 95,320 spectra took 40 minutes (when training for 15 iterations).

Spec2Vec is not meant to be the endpoint but rather a start of a new direction in spectral similarity scores. The low computational costs of Spec2Vec similarity scores make it possible to run extensive searches on very large library datasets. This makes it particularly suited to act as a pre-selection funnel that can easily be extended with a set of computationally more expensive similarity measures in the future. One could think of adding relevant mass differences as input to train the model, for example following the approach of Kreitzberg et al.^24^.

Finally, metabolomics is increasingly used as a tool to understand metabolic profiles and to perform integrative systems biology; furthermore, the use of MS/MS data has been promoted by community based platforms such as MassBank^5^, MS2LDA^23^ and GNPS^7^. We would like the community to use our novel score and therefore we created the modular matchms^25^ and Spec2Vec^26^ packages that can easily be incorporated in pipelines such as GNPS. It could also be used in other systems that rely upon spectral similarity. For example, SIRIUS and IOKR both use kernel matrices that are constructed from spectral similarities and whether Spec2Vec similarities could improve these pipelines is clearly worth investigation. In the present work, we have only demonstrated performance on LC-MS data. A very promising avenue will hence be to assess the utility of Spec2Vec for GC-MS data. When measuring molecules with GC-MS, precursor m/z values are usually not measured. This means that the initial precursor filtering to reduce the number of similarity calculations that can be done for LC-MS is not possible suggesting that the time savings available with Spec2Vec would be particularly desirable. Our work represents the first machine learning inspired spectral similarity score and, given its central place in metabolomics analyses, we believe Spec2Vec opens up new possibilities that will impact metabolomics analyses across all disciplines including clinical, food, and microbial metabolomics as well as biomarker and natural products discovery by improving identification, annotation, and networking.

## Methods

### Data preparation

The current study was done using a large LC-MS dataset provided on GNPS containing 154,820 spectra (https://gnps-external.ucsd.edu/gnpslibrary/ALL_GNPS.json from 2020-05-11, the raw data can be now be found on https://doi.org/10.5281/zenodo.3979010). The provided metadata was cleaned and corrected using matchms^25^ resulting in 94,121 spectra with InChIKey annotations. Using key metadata information such as compound names, chemical formulas and parent masses, an extensive automated lookup search was run against PubChem^27^ (via pubchempy^28^). As a result, 128,042 out of 154,820 spectra could be linked to an InChIKey (14,978 unique InChIKeys in the first 14 characters).

The here used **AllPositive** subset contains all spectra with positive ionization mode containing 112,956 spectra, out of which 92,954 with InChIKey (13,505 spectra with unique planar InChIKeys in first 14 characters before Spec2Vec related filtering).

We also worked with the considerably smaller subset **UniqueInchiKeys** which was reduced on purpose to be accessible for extensive benchmarking. It contains only one spectrum for every unique InChIKey from the **AllPositive** dataset (see supplemental or notebooks for details on the selection procedure).

In a next step we removed all peaks with m/z ratios outside the range [0, 1000] and discarded all spectra with less than 10 peaks. This left us with 95,320 spectra (out of which 77,092 with InChIKey) for the **AllPositive** dataset, and 12,797 spectra (all with InChIKey) for the **UniqueInchiKeys** dataset.

Since Spec2Vec similarity scores are conceptually very different from cosine-like similarity scores we decided to use two different peak filtering procedures for the two methods. For both the cosine and modified cosine score calculations we ignored all peaks with relative intensities <0.01 compared to the highest intensity peak. This is both to remove potential noise, but also to reduce the computational costs for the classical similarity scores.

Spec2Vec is comparing spectrum documents using language model analogies. For the underlying Word2Vec models we hence aimed at training on documents of comparable size which is achieved by removing excessive amounts of low intensity peaks. Since we expect that larger molecules on average will produce a higher number of meaningful fragmentation peaks, the maximum number of kept peaks per spectrum was set to scale linearly with the parent mass:

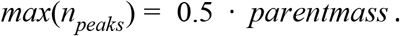

To assess if this procedure of having different peak filtering for the different similarity scores was indeed doing justice to the cosine and modified cosine score, we also repeated the library matching with cosine scores computed based on the Spec2Vec-processed data. This resulted in a slightly lower overall performance of the cosine score based library matching.

### From spectrum to document

After processing, spectra are converted to documents. For this, every peak is represented by a word that contains its position up to a defined decimal precision (““). For all presented results, a binning of two decimals was used, so that a peak at m/z 200.445 translates into the word “peak@200.45”. In addition to all peaks of a spectrum, neutral losses between 5.0 and 200.0 Da were added as. Neutral losses are calculated as *precursor_m/z_* − *peak_m/z_*. A list of all created peak and loss words is what we here refer to as a document.

A Word2Vec^20^ model is trained on all documents of a chosen dataset using gensim^29^. However, Spec2Vec in several aspects differs significantly from typical NLP applications so that some key parameters of the model also differ notably from the default settings. First of all, peaks in the mass spectra have no particular order that is comparable to the word order in a document. We hence set the window-size to 500, which in our case means that the entire spectrum (i.e. the entire document) counts as context. The two key types of word2vec models are skip gram and CBOW, the latter was generally observed to perform better in our case.

Although it is often considered that longer training will improve the results of a Word2Vec model, we found that this does not necessarily hold for our Spec2Vec spectral similarity measures. When using negative sampling during the training, model performance was observed to decrease for very long training runs. At the same time, however, including negative sampling allowed to obtain better overall results. Generally we found that training a model with negative sampling (negative=5) and 15 (**AllPositive**) up to 50 (**UniqueInchikey**) epochs were best suited for obtaining close to optimal model performance (see supplemental materials and fig. S4). To obtain a stable baseline performance we recommend to also train a model without negative sampling which will plateau for very long training runs.

### Spec2Vec similarity score

Spec2Vec similarity scores are derived on the basis of a pre-trained Word2Vec model. Word2Vec learns relationships between words (=peaks/losses) from co-occurrences across the seen documents. It then allows to represent words by abstract word vectors (so called word embeddings) in such a way that words of similar meaning are placed close to each other. With Spec2Vec our main interest lies in comparing entire spectra, which can be described by the sum of their words. To account for the dependency between peak relevance and peak intensity we calculate a spectrum vector *v_s_* as a weighted sum:

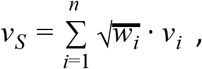

with *w_i_* the intensity (normalized to maximum intensity=1) and *v_i_* the word vector of peak *i*. For the similarity between two spectra we then compute the cosine score between two spectrum vectors:

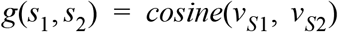

In practice we expect that Spec2Vec similarities will be most interesting to use when the underlying word2vec was trained on a large reference dataset containing many different fragments and losses and their structural relationships. It can also mean that it will be applied to spectra that were not part of the training data. In those cases some words (=peaks) of a given spectra might be unknown to the model. In those instances we can estimate the impact of the missing words by assessing the uncovered weighted part of a spectrum:

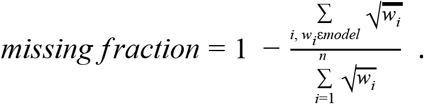

Having few unknown peaks of low intensity in a spectrum will count only little to the missing fraction, whereas high numbers of unknown peaks or few unknown peaks of high intensity will result in a high missing fraction. By setting a threshold for the missing fraction (e.g. <0.05), returning Spec2Vec similarity scores for spectra far outside the learned peaks (and losses) can be avoided.

### Structural similarity

Assessing the structural similarity between two molecules remains a complex topic. Finding the best measure to define structural similarity between two molecules lies outside the scope of this study; most recent studies converge to the Tanimoto similarity as one of the most practical and well-performing measures^30^. Thus, for the presented results, the structural similarity was measured by taking the Tanimoto similarity (Jaccard index) based on daylight-like molecular fingerprints (rdkit molecular fingerprints, version 2020.03.2, 2048 bits, derived using rdkit ^31^ via matchms^25^).

## Acknowledgement

We want to thank Marnix Medema and Michelle Schorn for many helpful discussions. Further, we thank the eScience Center generalization team for help and input around code design, packaging, and testing. J.J.J.v.d.H. acknowledges funding from an ASDI eScience grant, ASDI.2017.030, from the Netherlands eScience Center—NLeSC.

## Code, trained models, and data

The underlying code was developed into two Python packages to handle and compare mass spectra, matchms (https://github.com/matchms/matchms) and spec2vec (https://github.com/iomega/spec2vec). Both packages are freely available and can be installed via conda ^25,26^.

All additional functions to analyse the data and to create the presented plots can be found under https://github.com/iomega/spec2vec_gnps_data_analysis. This repository also contains extensive Jupyter notebooks to document the entire workflow from raw data to the figures presented in this work. The notebooks were tested using matchms 0.6.0 and spec2vec 0.3.1. The two most important trained Word2Vec models used in this work can be downloaded from https://doi.org/10.5281/zenodo.3978054 (trained on **UniqueInchikey** dataset) and https://doi.org/10.5281/zenodo.3978070 (trained on AllPositive dataset).

The pre-processed, cleaned dataset with all positive ionization mode spectra can be downloaded from https://doi.org/10.5281/zenodo.3978118, the original raw data can be accessed from https://doi.org/10.5281/zenodo.3979010.

Calculated all-vs-all similarity score matrices for cosine score, modified cosine score, and fingerprint-based similarity (Tanimoto) for the **UniqueInchikey** dataset can be found on https://zenodo.org/record/3979074.

## Notes

https://zenodo.org/record/3978118

### Competing Interest Statement

The authors have declared no competing interest.

### Summary of Updates

Revised figure 1,2,3,5 to make them more easily understandable.Numerous minor text edits to enhance readability. Proper split into main text and supplemental. Refer to used matchms and spec2vec versions. Added authors.

