## Supplemental Material for "Spec2Vec: Improved mass spectral similarity scoring through learning of structural relationships"

Affiliations:


#### Content:


| | |
|---|---|
| Data processing protocol | 1 |
| Analysis of modified cosine reliability | 3 |
| Benchmarking and parameter tuning for the cosines score | 5 |
| Benchmarking and parameter tuning for the modified cosines score | 9 |
| Spec2Vec model parameters | 11 |
| Network analysis | 12 |


#### Data processing protocol

**Data pre-processing:**

1. Import all spectra from GNPS website via matchms json import (all spectra file from https://gnps-external.ucsd.edu/gnpslibrary/ALL_GNPS.json from 2020-05-11)
   Found non-empy spectra: 154820 spectra
2. Run default matchms filters to clean, correct, and infer missing metadata. Results in:
   94462 spectra with InChI (15822 unique)
   94155 spectra with Smiles (20542 unique)
   94121 spectra with InChIKey (13505 unique in first 14 characters)

3. Run extensive automated PubChem lookup (code provided in manuscript repository).
   Results in:
   128103 spectra with InChI (17620 unique)
   128052 spectra with Smiles (23097 unique)
   128042 spectra with InChIKey (14978 unique in first 14 characters)

**Creation of subsets:**

Creation of 4 different datasets:

- **AllGNPS:** All spectra after matchms cleaning and PubChem lookups (154820 spectra)
- **AllPositive:** all spectra from AllGNPS dataset with positive ionization mode.
  Total: 112956 spectra
  92997 spectra with InChI (16071 unique)
  92964 spectra with Smiles (20540 unique)
  92954 spectra with InChIKey (13717 unique in first 14 characters)
- **AllPositiveAnnotated**
- **UniqueInchiKeys:** Reduced dataset used for benchmarking. Keep only one spectrum for every unique InChIKey from the AllPositive dataset. Spectra are selected by:
  i) If possible, select spectra with > 10 peaks above 1% of maximum peak intensity
  ii) Out of those (if multiple): select best library quality level: 1> 2> 3
  iii) Out of those (if multiple): select spectrum with most peaks above 1% of maximum peak intensity
  iv) if still multiple, pick random!
  This gives 13717 spectra, out of which 12797 spectra contain >= 10 peaks.

**Data post-processing for "classical" similarity scores (cosine, modified cosine)**

1. Remove peaks with m/z outside [0, 1000]
2. Remove spectra with < 10 peaks remaining
3. Remove peaks with intensities < 0.01 maximum peak intensity (smaller peaks will slow down score calculation, but won't contribute much to the overall scores)

**Data post-processing for spec2vec**

1. Remove peaks with m/z outside [0, 1000]
2. Remove spectra with < 10 peaks remaining
3. Reduce number of peaks using matchms "reduce_number_of_peaks" filter.
   Parameters: n_required = 10, ratio_desired = 0.5
   Spec2vec is comparing spectrum documents using language model analogies. For the underlying word2vec models we aimed at training on documents of roughly comparable size to ensure that spectra will also get comparable attention during model training. The raw data contained spectra with numbers of peaks ranging from 10 (our own set

threshold to include spectra) up to several 10,000s of peaks. We hence removed excessive amounts of low intensity peaks. To account for the fact that larger molecules on average show more fragmentation peaks, the maximum number of kept peaks per spectrum was set to scale linearly with the parent mass:

$$max(n_{peaks}) = 0.5 \cdot parentmass .$$

#### Analysis of modified cosine reliability

We noted a large amount of false positive among the spectra pairs with high cosine or modified cosine scores. With false positives we here mean spectral pairs that receive a high spectra similarity but a low structural similarity score. This clearly shows in histograms of all structural similarities which belonged to pairs that received spectra similarity scores above a set threshold. For a given score, for instance modified cosine, we would select all pairs across the **UniqueInchikey** dataset (12,797 spectra, total of 81,875,206 unique pairs when excluding pairs of spectra with themselves) with a modified cosine score > 0.7, 0.8, etc. and create a histogram of the corresponding structural similarity values (fig. S1). Even though we already use a relatively small peak m/z tolerance of 0.005 Da (preventing the collapse of too many mass fragments with different elemental formulas into one mass bin), the *min_match* criteria also clearly has to be raised to obtain acceptable results. In comparison, Spec2Vec similarities show a visibly lower amount of false positives (Fig. S1, lower right).

We also consider this one of the reasons for the notably poorer correlation between cosine and modified cosines scores and structural similarities (Figure 3 in main manuscript).

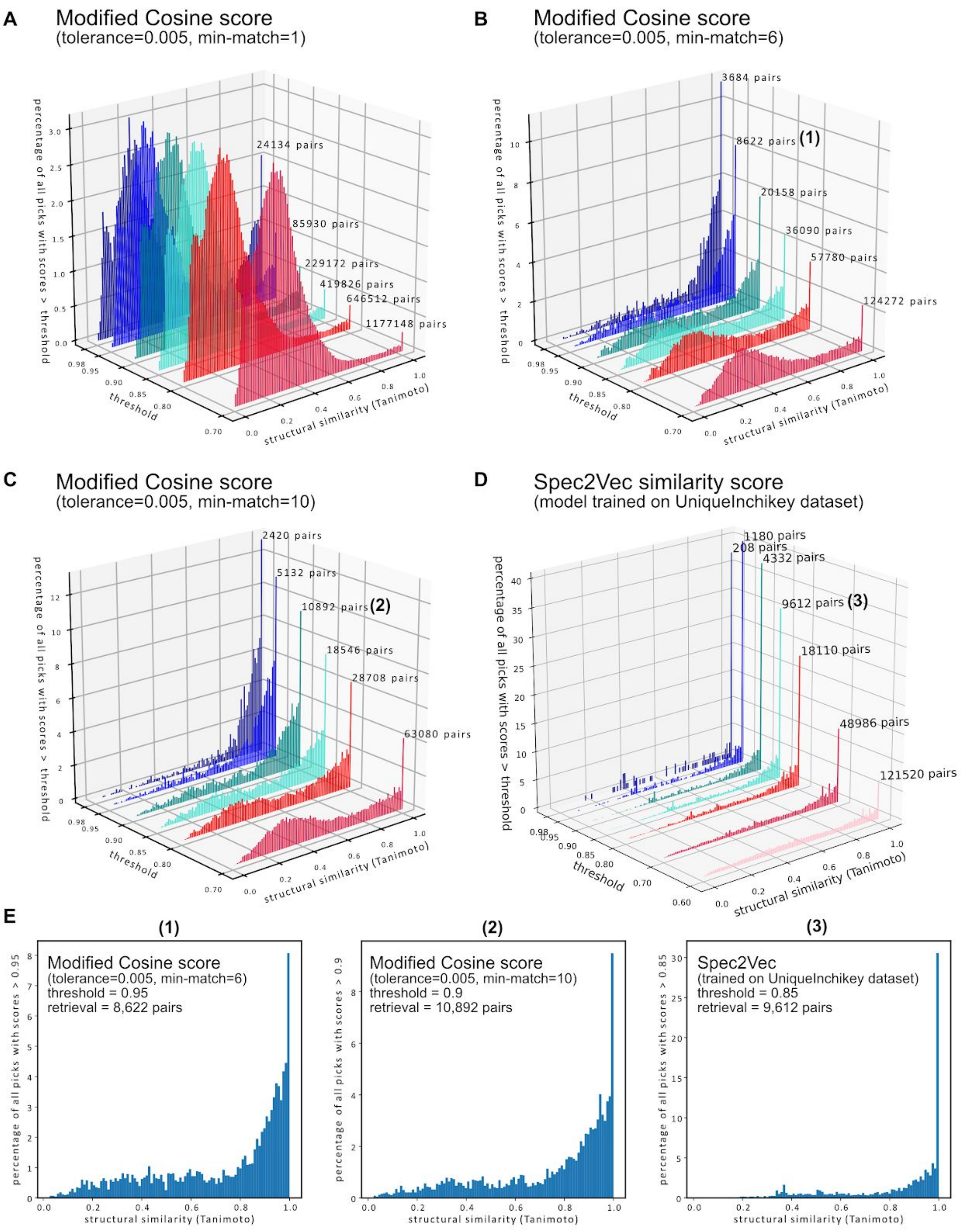


**Figure S1**. Histograms of all structural similarity scores (Tanimoto) within the UniqueInchikey dataset for which the respective similarity scores are above a set threshold. Scores between identical spectra are excluded here. Thresholds between 0.7 to 0.98 were used to create the displayed histograms and are shown on the plot axis. The number of pairs that fitted the criteria (min-match and > threshold) is written on the top right of each histogram. (A) Modified cosine scores without requiring multiple machine peaks between two spectra results in drastic numbers of false positives (=low Tanimoto scores despite spectra similarity scores > threshold). (B) Raising the minimum number of matches to 6 already improves the reliability of the modified cosine score notably. Still, even for comparably high thresholds (scores > 0.9 or > 0.95) a considerable fraction of all pairs with high modified cosine scores do not correspond to similar molecules. (C) For a minimum of 10 matching peaks the reliability further improved, but also the number of pairs fulfilling this criteria continues to drop. (D) Spec2Vec similarities also show notable levels of false positives, in particular for lower thresholds (0.7), but generally are visibly more shifted towards high Tanimoto scores when compared to the modified Cosine scores. Absolute threshold values are hard to compare. For instance, a Spec2Vec similarity of 0.9 could be more or less common than a Cosine score of 0.9 for a given setting. In (E) we hence compare three histograms that represent roughly equal numbers of spectra pairs (between 8,622 and 10,892). This shows that for comparable retrieval rates, high Spec2Vec scores correlate less frequently with very low molecular similarities.

#### Benchmarking and parameter tuning for the cosines score

Cosine scores were calculated for all-vs-all spectra of the **UniqueInchikey** dataset using different tolerances and minimum matching peak thresholds. This required computing similarity scores for all possible pairs between the 12,797 spectra (81,875,206 unique pairs), which --depending on the pre-processing and parameters used-- takes several hours on a standard CPU to compute. The precise computation time is highly dependent on the peak filtering, for the present settings, run on a Intel i7-8550U, the computation took about 6 hours. The number of matching peaks and the actual scores can be computed simultaneously (using matchms[25]).

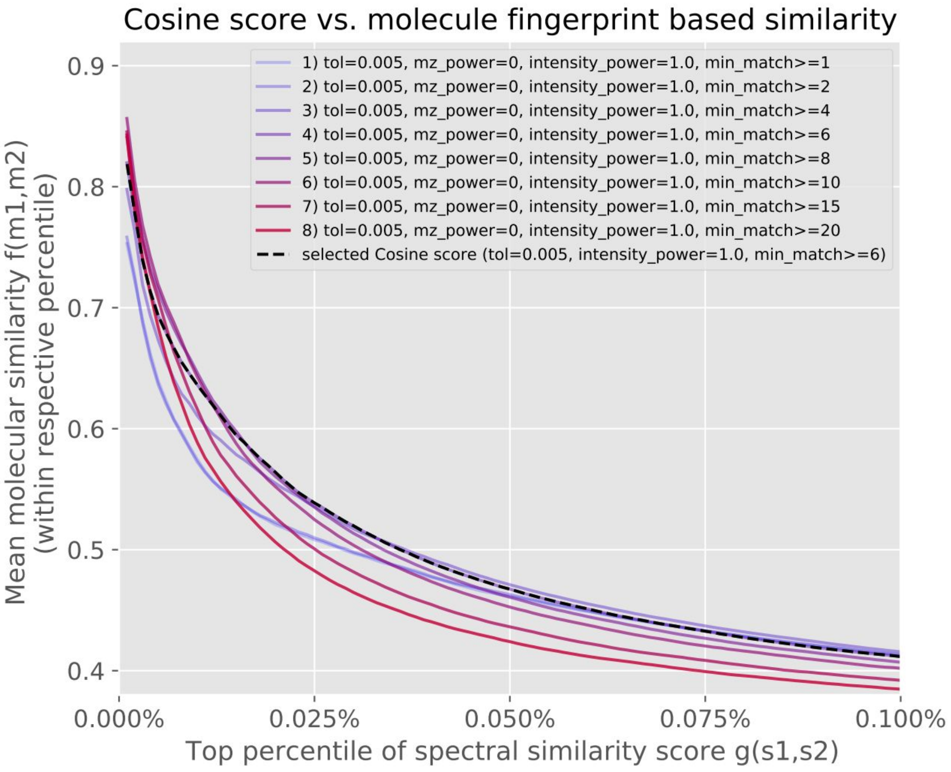


**Figure S2**. Parameter search for cosine similarity score. Varying from 1 to 20 the minimum number of matching peaks necessary to calculate a cosine score. For all following comparisons, min_match=6 was chosen.

Cosine scores come in many flavors, which makes it impossible to systematically compare all possible parameters settings and implementations. Apart from what we would consider the most basic implementation used here, we also tested cosine scores that weigh peaks according to their m/z ratio or their relative intensity. Settings that were tested in-depth stem from a suggestion from Demuth et al.[11] , as well as from current implementation used in NIST and Massbank[12].

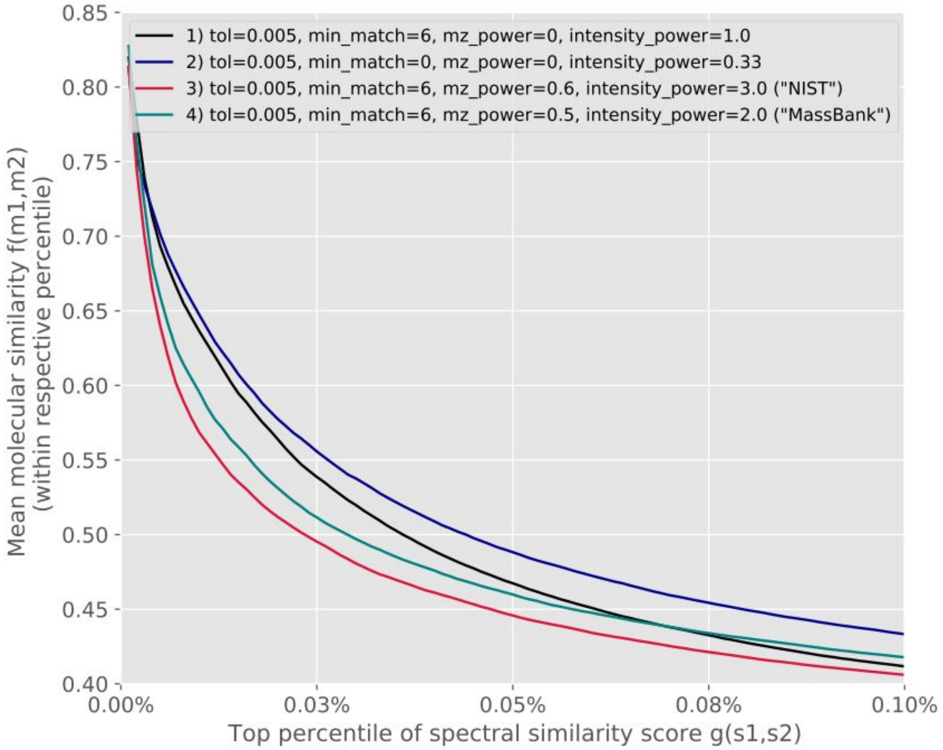


**Figure S3**. Comparing different Cosine score flavors. Four different mz_power/intensity_power parameter settings were tested. The plot shows the respective results with the best performing min_match criterion.

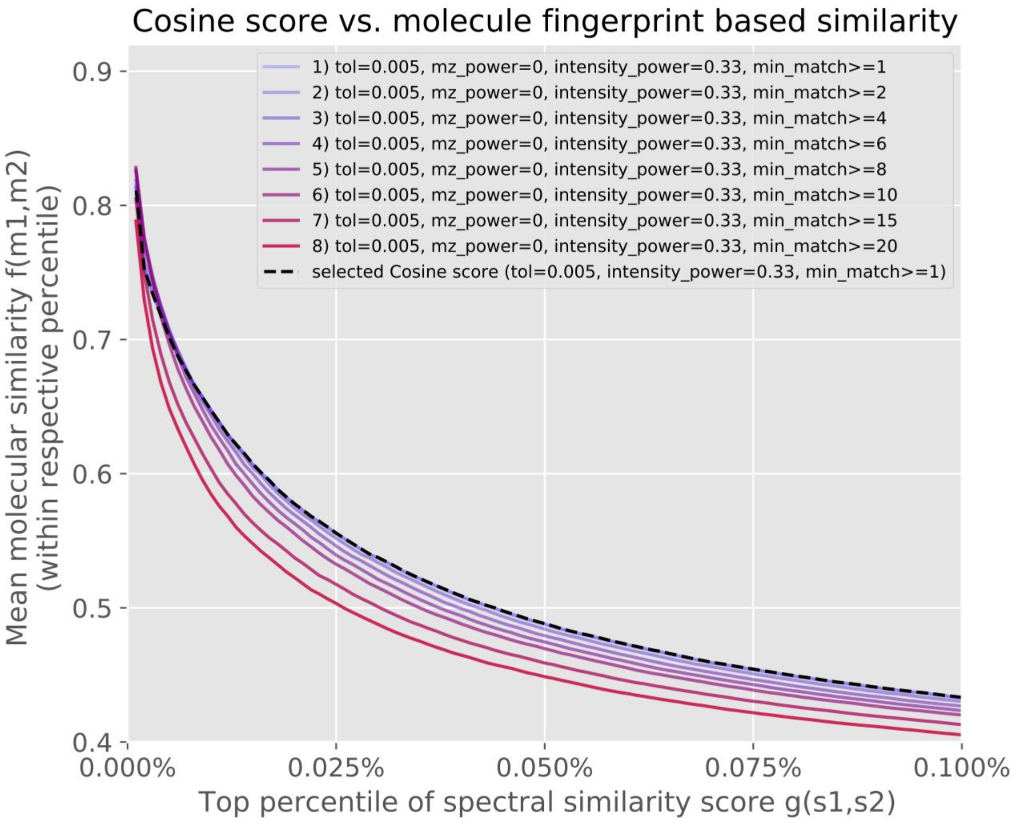


**Figure S4**. Benchmarking of cosine score with intensity_power=0.33 (Demuth et al. [11]) for different min_match settings.

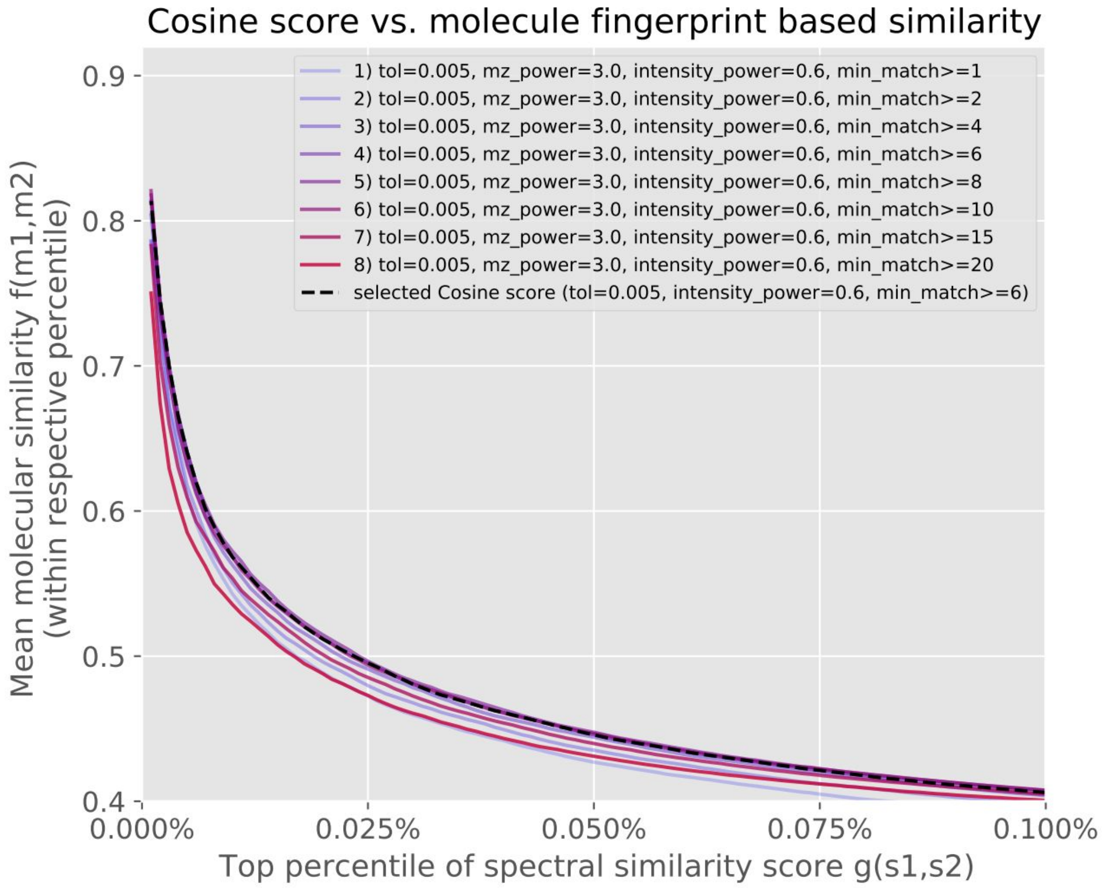


**Figure S5**. Benchmarking of cosine score with mz_power=3.0 and intensity_power=0.6 (NIST settings [12]) for different min_match settings.

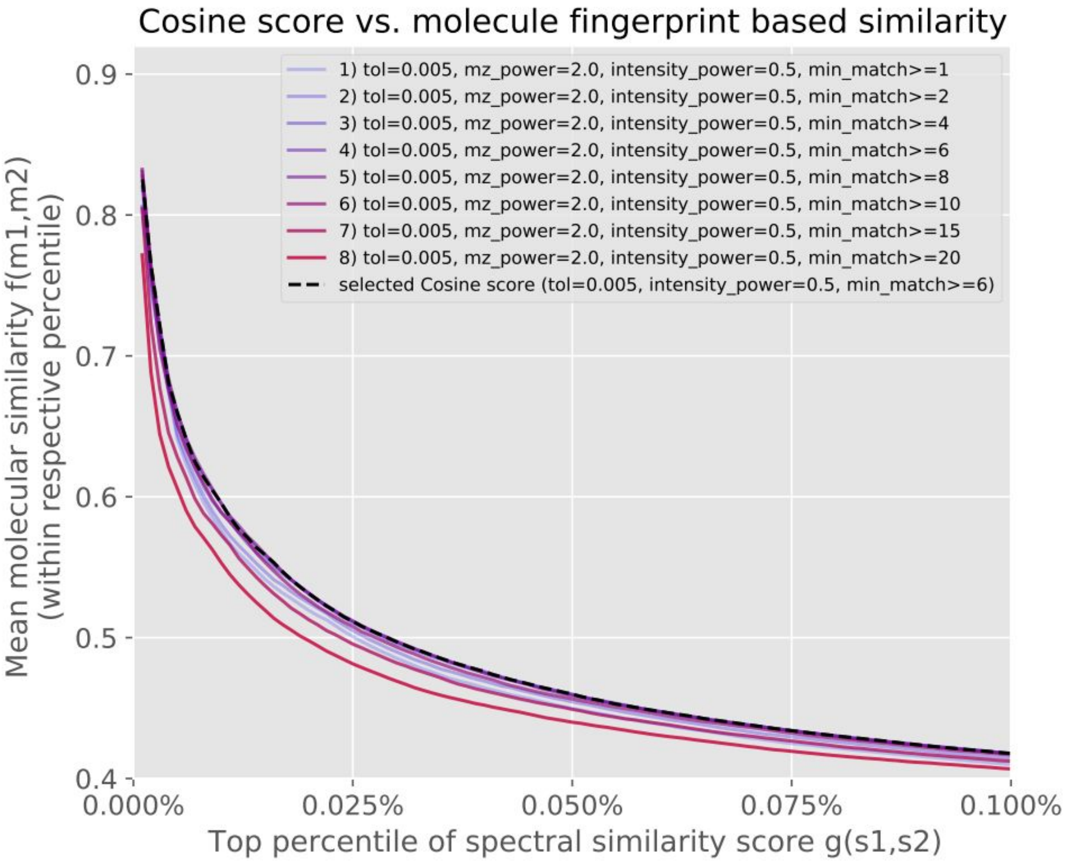


**Figure S6**. Benchmarking of cosine score with mz_power=2.0 and intensity_power=0.5 (MassBank settings [12]) for different min_match settings.

### Benchmarking and parameter tuning for the modified cosines score

As for the cosine score, modified cosine scores were calculated for all-vs-all spectra of the **UniqueInchikeys** dataset using a tolerance of 0.005 Da together with different minimum matching peak thresholds (fig. S7).

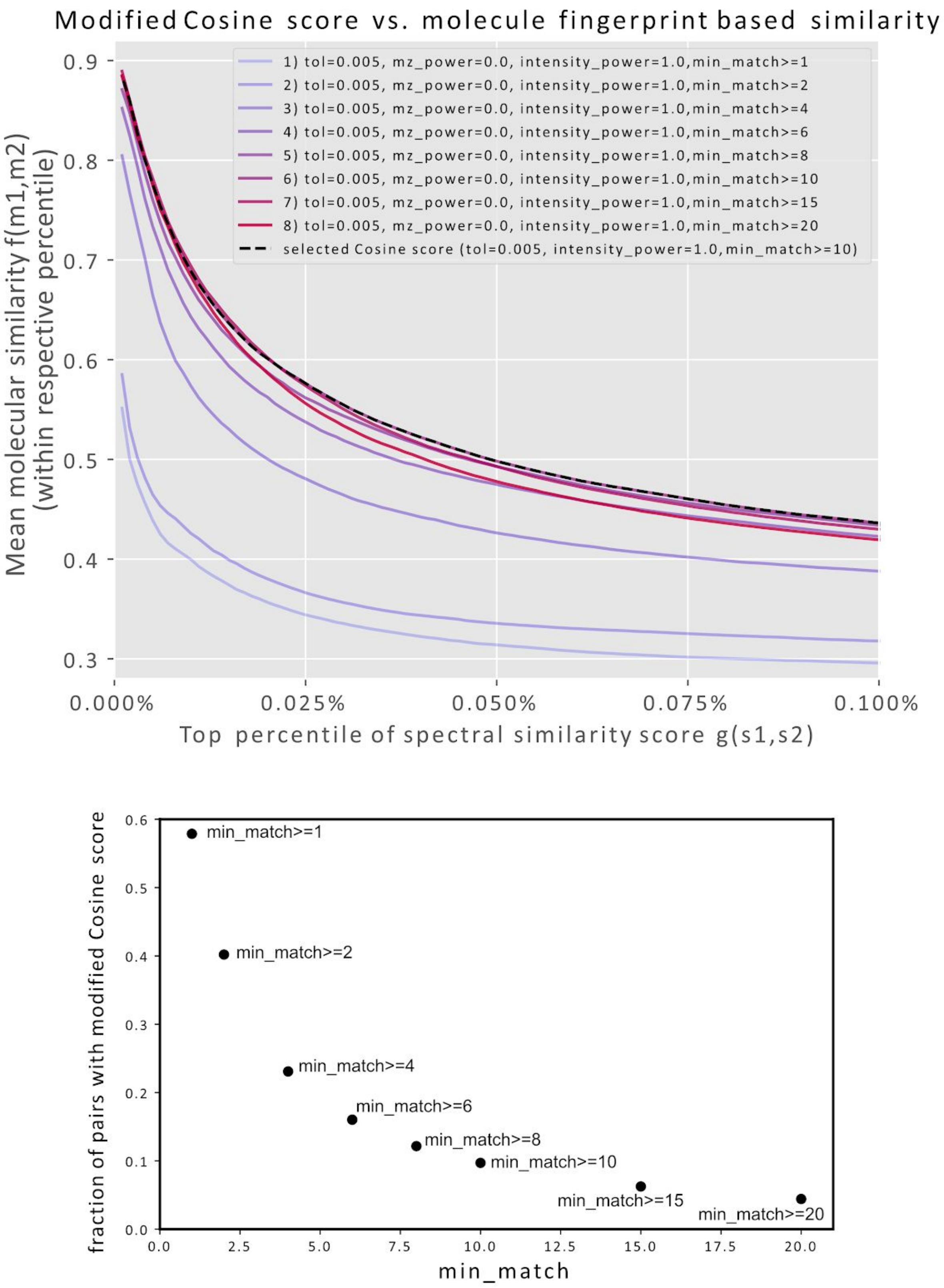


**Figure S7**. Parameter search for modified cosine similarity score. Varying from 1 to 20 the minimum number of matching peaks necessary to calculate a cosine score. For all following comparisons, min_match=10 was chosen. Please note that with increasing min_match parameter, more and more spectra pairs will not receive a Cosine score (bottom plot). For min_match=10, for instance, less than 10% of all spectra pairs will receive a Cosine score. For min_match=20 this drops below 5%. This also means potentially losing correctly matching pairs that just happen to not have sufficient shared peaks (unlike for Spec2Vec which would return a score for any pair).mat

### Spec2Vec model parameters

The underlying word2vec models were trained using the spec2vec Python library running Gensim. The key model parameters were

- Window size = 500
- Word vector dimension = 300
- Negative sampling
- CBOW mode
- Initial learning rate = 0.025
- Learning rate decay per iteration (per epoch) = 0.00025

We compared models after different iterations using the benchmarking dataset **UniqueInchikey** to find suitable numbers of training iterations (fig. S8). Generally training was comparably stable over a larger range of iterations. It converges to stable results when switching off negative sampling, but negative sampling resulted in better overall results (not shown here).

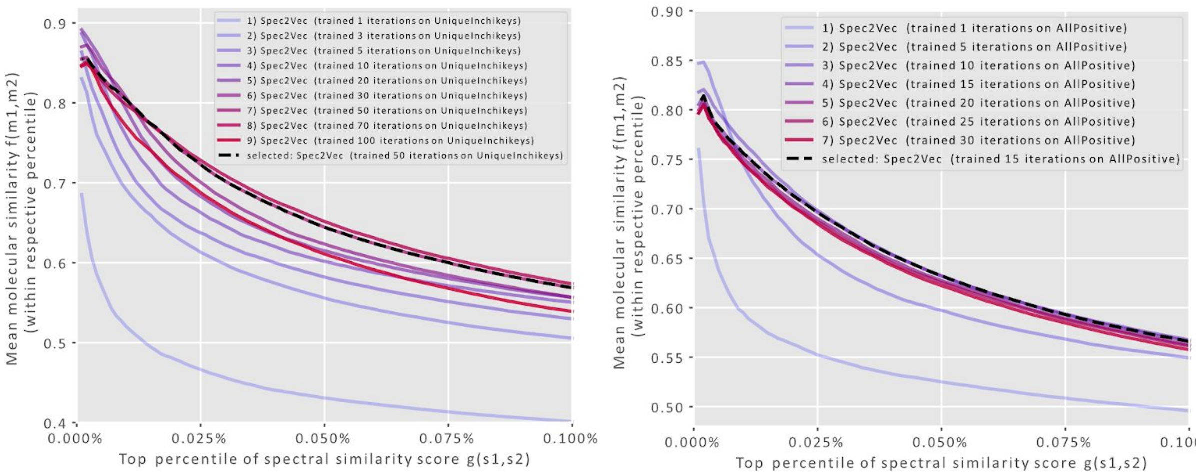


**Figure S8.** Word2Vec models were trained on both the **UniqueInchikey** and the **AllPositive** dataset over many iterations. The overall performance was monitored by the correlation between Spec2Vec similarities and structural similarity and revealed that the model rapidly improved during the first 10-15 epochs (iterations). For the **UniqueInchikey** dataset we found the best performance around 50 iterations, for the AllPositive dataset we observed that the changes after 15 iterations were rather minor.

### Network analysis

Based on the different working principles of cosine-based spectral similarity scores, and Spec2Vec similarity, we expect different weaknesses and strengths of both score types. Those differences could be exploited by combining scores. In a first, and highly simplified, test on the described molecular networking task (see main manuscript, fig.6), we observed that a simple linear combination of both modified cosine and Spec2Vec similarities only mildly lowers the total fraction of clustered spectra, but notably reduces fraction of poorly clustered spectra (fig. S9). We can hence increase clustering accuracy by relying on both scores, which seems a promising starting point to build upon in future work.

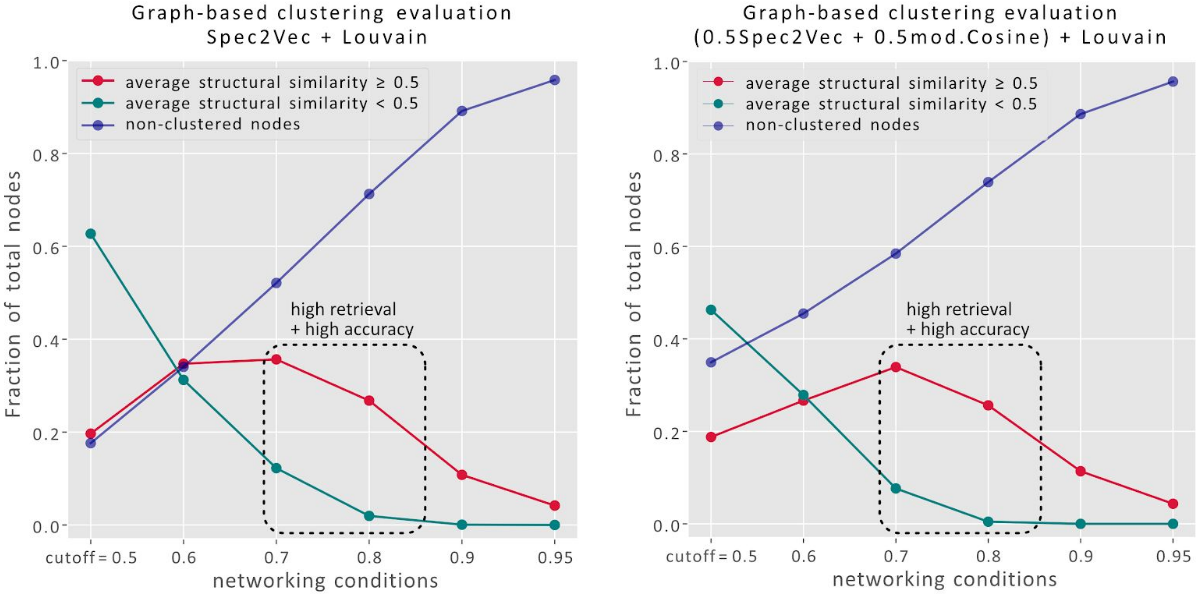


**Figure S9.** Networks were generated from spectra (nodes) by adding links based on spectra similarities, as shown in the main manuscript (see fig. 6). The left plot is identical to the right plot in figure 6 (main manuscript) and was generated by only using Spec2Vec similarity scores. On the right we used a simple combination of Spec2Vec similarities and modified Cosine scores (similarity = 0.5 Spec2Vec similarity + 0.5 mod.Cosine, min_match=10, tolerance=0.005). Already this simple combination leads to a drop of the loss structural similarity clusters (green) with respect to the well-clustered fraction (red), effectively leading to a higher accuracy. We expect that more refined combinations of both similarity scores can be used to further build upon these results.
